# MrTADFinder: A network modularity based approach to identify topologically associating domains in multiple resolutions

**DOI:** 10.1101/097345

**Authors:** Koon-Kiu Yan, Shaoke Lou, Mark Gerstein

## Abstract

Genome-wide proximity ligation based assays such as Hi-C have revealed that eukaryotic genomes are organized into structural units called topologically associating domains (TADs). From a visual examination of the chromosomal contact map, however, it is clear that the organization of the domains is not simple or obvious. Instead, TADs exhibit various length scales and, in many cases, a nested arrangement. Here, by exploiting the resemblance between TADs in a chromosomal contact map and densely connected modules in a network, we formulate TAD identification as an optimization problem and propose an algorithm, MrTADFinder, to identify TADs from intra-chromosomal contact maps. MrTADFinder is based on the network-science concept of modularity. A key component of it is deriving an appropriate background model for contacts in a random chain, by numerically solving a set of matrix equations. The background model preserves the observed coverage of each genomic bin as well as the distance dependence of the contact frequency for any pair of bins exhibited by the empirical map. Also, by introducing a tunable resolution parameter, MrTADFinder provides a self-consistent approach for identifying TADs at different length scales, hence the acronym “Mr” standing for Multiple Resolutions. We then apply MrTADFinder to various Hi-C datasets. The identified domains are marked by boundary signatures in chromatin marks and transcription factor (TF) that are consistent with earlier work. Moreover, by calling TADs at different length scales, we observe that boundary signatures change with resolution, with different chromatin features having different characteristic length scales. Furthermore, we report an enrichment of HOT regions near TAD boundaries and investigate the role of different TFs in determining boundaries at various resolutions. To further explore the interplay between TADs and epigenetic marks, we examine how somatic mutations are distributed across boundaries (as tumor mutational burden is known to be coupled to chromatin structure), finding a clear stepwise pattern. Overall, MrTADFinder provides a novel computational framework to explore the multi-scale structures in Hi-C contact maps.

**Author Summary:** The accommodation of the roughly 2m of DNA in the nuclei of mammalian cells results in an intricate structure, in which the topologically associating domains (TADs) formed by densely interacting genomic regions emerge as a fundamental structural unit. Identification of TADs is essential for understanding the role of 3D genome organization in gene regulation. By viewing the chromosomal contact map as a network, TADs correspond to the densely connected regions in the network. Motivated by this mapping, we propose a novel method, MrTADFinder, to identify TADs based on the concept of modularity in network science. Using MrTADFinder, we identify domains at various resolutions, and further explore the interplay between domains and other chromatin features like transcription factors binding and histone modifications at different resolutions. Overall, MrTADFinder provides a new computational framework to investigate the multiple length scales that are built inside the organization of the genome.

## Introduction

The packing of a linear eukaryotic genome within a cell nucleus is dense and highly organized. Understanding the role of 3D genome in gene regulation is a major area of research [1][2][3][4]. Recently, genome-wide proximity ligation based assays such as Hi-C have provided insights into the complex structure by revealing various structural features regarding how a genome is organized [5][6][7]. Perhaps, one of the most important discoveries is the domain of self-interacting chromatin called topologically associating domain (TAD) [8][9]. Inside a TAD, genomic loci interact often; but between TADs, interactions are less frequent. Thus the TAD emerges as a fundamental structural unit of chromatin organization; it plays a significant role in mediating enhancer-promoter contacts and thus gene expression, and breaking or disruption of TADs can lead to diseases like cancers [10][11][12]. Therefore, a deeper understanding of TADs from Hi-C data presents an important computational problem.

Results of a typical Hi-C experiment are usually summarized by a so-called chromosomal contact map [5]. By binning the genome into equally sized bins, the contact map is essentially a matrix whose element (*i*, *j*) reflects the population-averaged co-location frequencies of genomic loci originated from bins *i* and *j*. In this representation, TADs are displayed as blocks along the diagonal of a contact map [8][9]. Despite the fact that TADs are rather eye-catching in a contact map, computational identification is still challenging because of experimental factors such as noise and inadequate coverage. Moreover, it is apparent from a visual examination of the contact map that TADs exhibit various length scales: there are TADs that appear to be overlapping, and within many TADs, there are rich sub-structures.

Mathematically speaking, it is very natural to transform a contact matrix to a weighted network in which nodes are the genomic loci (or bins) whereas the interaction between two loci is quantified by a weighted edge. In network science, a widely studied problem is the identification of network modules, also known as community detection problem [13]. A module refers to a set of nodes that are densely connected. In its simplest form, the community detection problem concerns with whether nodes of a given network can be divided into groups such that connections within groups are relatively dense while those between groups are sparse. Therefore, by viewing the chromatin interactions as a network, the highly spatially localized TADs immediately resemble densely connected modules. Motivated by the resemblance, we formulate the identification of TADs as a global optimization problem based on the observational contact map and a background model. As a network-based approach, our method goes beyond a direct adaptation of standard community detection algorithms. We introduce a novel background model that takes into account the effect of genomic distance, which is specific to the context of genome organization. The objective function is optimized using a heuristic algorithm that is efficient even if the size of the input contact map is large. Furthermore, by introducing a tuning parameter, our network approach can identify TADs at different resolutions. At a low resolution, larger TADs are found whereas, at a high resolution, smaller TADs are identified as the nucleome is viewed on a finer scale. In other words, the method can identify TADs at different length scales. We name our method MrTADFinder where the acronym Mr stands for multiple resolutions.

## Results

### A network modularity framework for TADs identification

The identification of modules in a network is formulated as a global optimization problem on the so-called modularity function over possible divisions of the network. Consider an unweighted network represented by an adjacency matrix *A*. For a particular division (i.e. a mapping from the set of all nodes to a set of modules), the modularity is defined as the fraction of edges within modules minus the expected fraction of such edges in a randomized null model of the network. Mathematically, the modularity is equal to

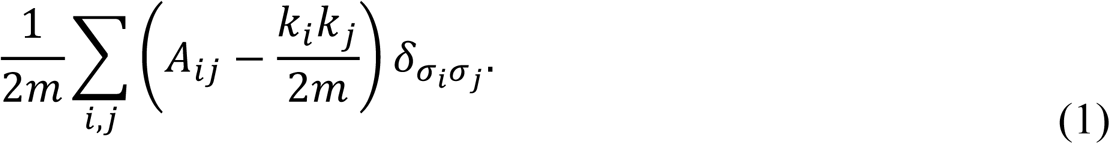

Here, the summation goes over all possible pairs of nodes, the value of the Kronecker data *δ*_*σiσj*_ equals one if nodes *i* and *j* have the same label σ and zero otherwise, meaning only pairs of nodes within the same module are summed. In particular, *m* is the number of edges in the network whereas the expression *k*_*i*_*k*_*j*_/2*m* represents the expected number of edges between *i* and *j* in a so-called configuration model. The configuration model is a randomized null model in which the degrees of nodes *k*_*i*_ are fixed to match those of the observed network, but edges are in other respects placed at random. High values of the modularity correspond to good partitions of a network into modules and similarly low values to bad partitions. Optimizing the modularity function leads us to the best partition over all possible partitions. More recently, a so-called resolution parameter *γ* has been incorporated in equation (1) to adjust the size of the resultant modules [14].

Following the network formalism, given a Hi-C contact map represented by a weighted matrix *W*, we define a similar objective function *Q* as

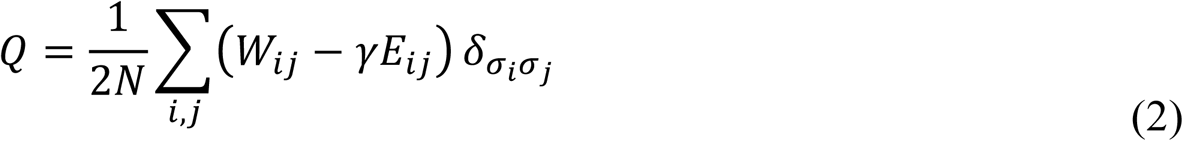

Here, *i*, *j* index the equally binned genomic loci. *N* is the total number of pair-end reads. *E*_*ij*_ is the expected number of contacts between locus *i* and locus *j*. *γ* is the resolution parameter that could be used to tune the size of resultant TADs. Very much similar to the network setting, the identification of TADs aims to partition the loci into domains such that *Q* is optimized. Nevertheless, it is important to emphasize two points. First, unlike the case in a network, the bins in a chromosome form a continuous chain and therefore genomic loci belonging to a TAD have to form a continuous segment. Second, simply because of the physical nature of chromosome, the expected number of contacts between locus *i* and locus *j* depends on their genomic distance. Two loci that are close together in a 1-dimensional sense are expected to have a higher contact frequency as compared to two loci that are far apart. This point suggests that the null model *E*_*ij*_ in equation (2) has to be modified.

### A novel null model of intra-chromosomal contact maps

Thus, given an intra-chromosomal contact map *W*, the expected null model E is defined as

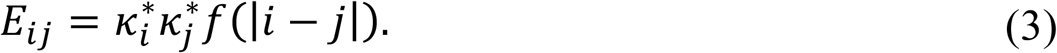

Here, *f* is the average number of contacts as a function of distance *d* = |*i* − *j|*. By considering all possible pairs of bins in *W* in terms of their distance apart and the contact frequency, we estimate *f* by local smoothing (see Methods). For intermediate values of *d*, *f* follows pretty well with a power-law function d^-1^ (see Figure S1), which is a well-known observation first reported in [5].

As a null model, the resultant *E* matrix satisfies a set of constraints, namely

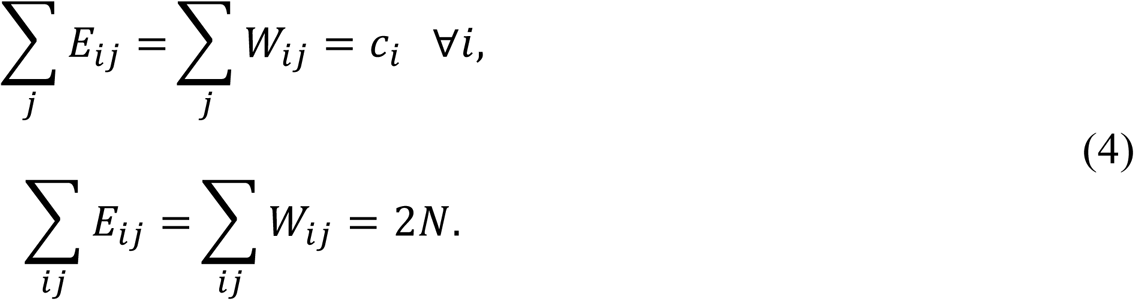

The first equation means that the coverage *c*_*i*_, i.e. the total number of reads (one end of pair-end reads) mapped to bin *i*, defined in the observed map is the same as the coverage defined in the null model. The second equation is a direct consequence of the first equation, where *N* is the total number of pair-end reads mapped to the chromosome. As *f* has been estimated from the observed *W*, we can numerically solve all the unknowns 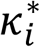 in the system of matrix equations (see Methods). Mathematically, 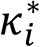 can be regarded as an effective coverage because of the correlation between 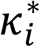 and the coverage *c*_*i*_ is extremely high (r=0.95, Figure S2). In comparison with equation (1), 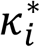 is conceptually analogous to the degree *k_i_*. As shown in Figure 1, given a particular matrix *W*, the contact frequency of the resultant null model *E* are the highest in the diagonal and decrease gradually away from the diagonal. With *W* and *E,* for any given resolution parameter *γ*, we employ a modified Louvain algorithm to optimize *Q* (see Methods and Figure 1 for details). To ensure robustness, multiple runs of the modified Louvain algorithm are performed, and a boundary score is defined as the fraction of times a bin is called as a boundary. The final set of TADs is defined based on the set of consensus boundaries (Figure 1 and Methods). It is important to emphasize that the conventional Louvain algorithm used in network analysis [15] cannot be directly used because chromatin domains are continuous segments.

**Figure 1:**
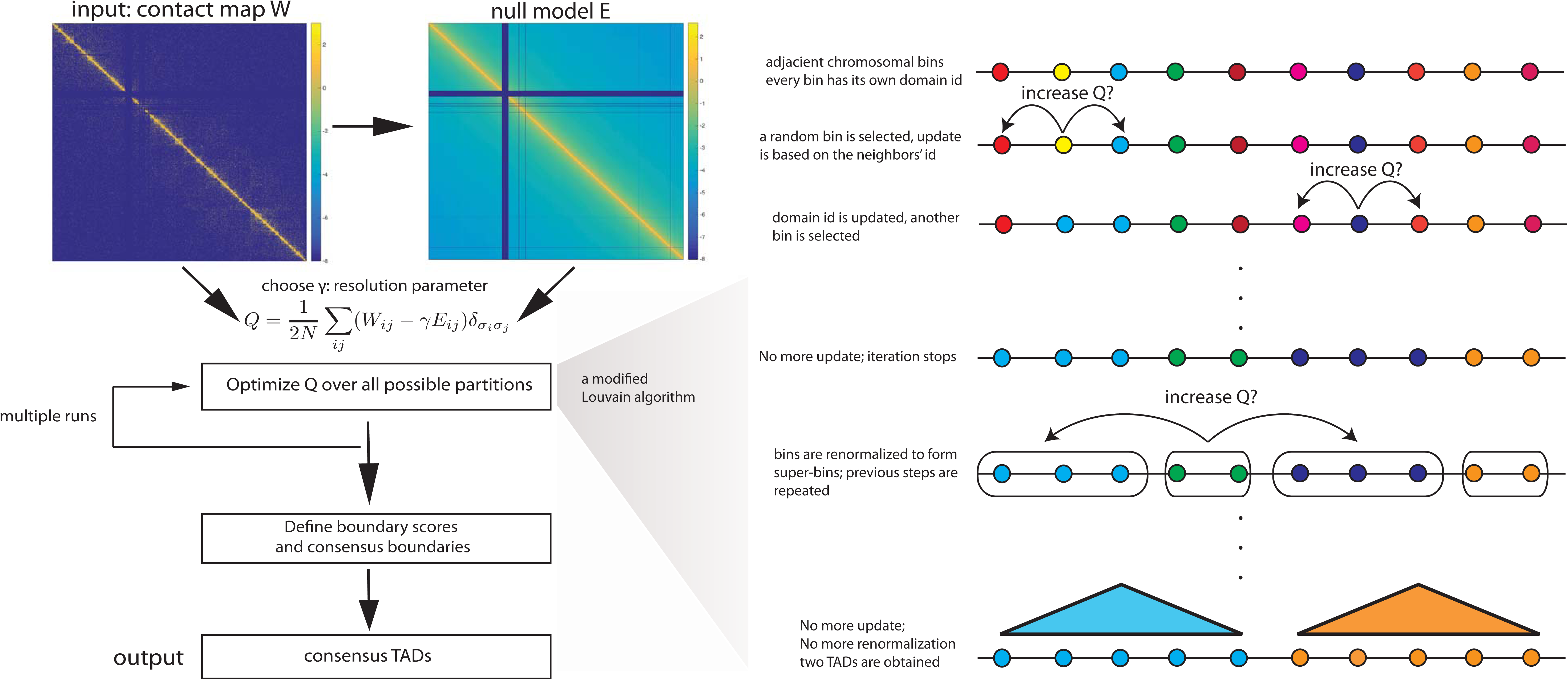
Overview of MrTADFinder. The input of MrTADFinder is an intra-chromosomal contact map *W*. A null model *E* is obtained from W. Given a particular resolution γ; the chromosome is partitioned probabilistically in a way such that the objective function Q is maximized. The optimization is performed by a modification Louvain algorithm shown on the right. The algorithm is stochastic because the updating order of nodes is random. A boundary score is defined after multiple trials for all adjacent bins. Adjacent bins that are robustly assigned to two different TADs form a consensus boundary. The output of MrTADFinder is a set of consensus domains bound by the consensus domains.

### Identifying TADs in multiple resolutions

As a demonstration, we applied MrTADFinder to analyze Hi-C data of hES cell from [8]. Figure 2A shows a particular snapshot of the contact map (for chromosome 10) and its alignment with the identified TADs. In general, the TADs displayed agree well with the apparent block structures in the contact map. Of particular interest is the choice of *γ* that capture various length scales in domain organization. As shown in Figure 2A, when *γ* increases, a large TAD breaks into a few small TADs. On the other hand, a few large TADs merge together to form an even larger TAD as the value of *γ* is lowered. Statistically speaking, *γ* quantifies to what extent do we accept the enrichment of empirical contact frequency over the expectation. As *γ* increases, only matrix elements close to the diagonal contribute positively to the objective function. Therefore, in general, the size of TADs decreases (see Figure 2B) and the number of TADs increases (see Figure 2C). For example, when *γ*=1.0, there are about 1000 TADs in hES cells with a median size of 3Mb. When *γ*=2.25, the number of TADs increases to 2600 and the median size is roughly 1Mb.

**Figure 2.**
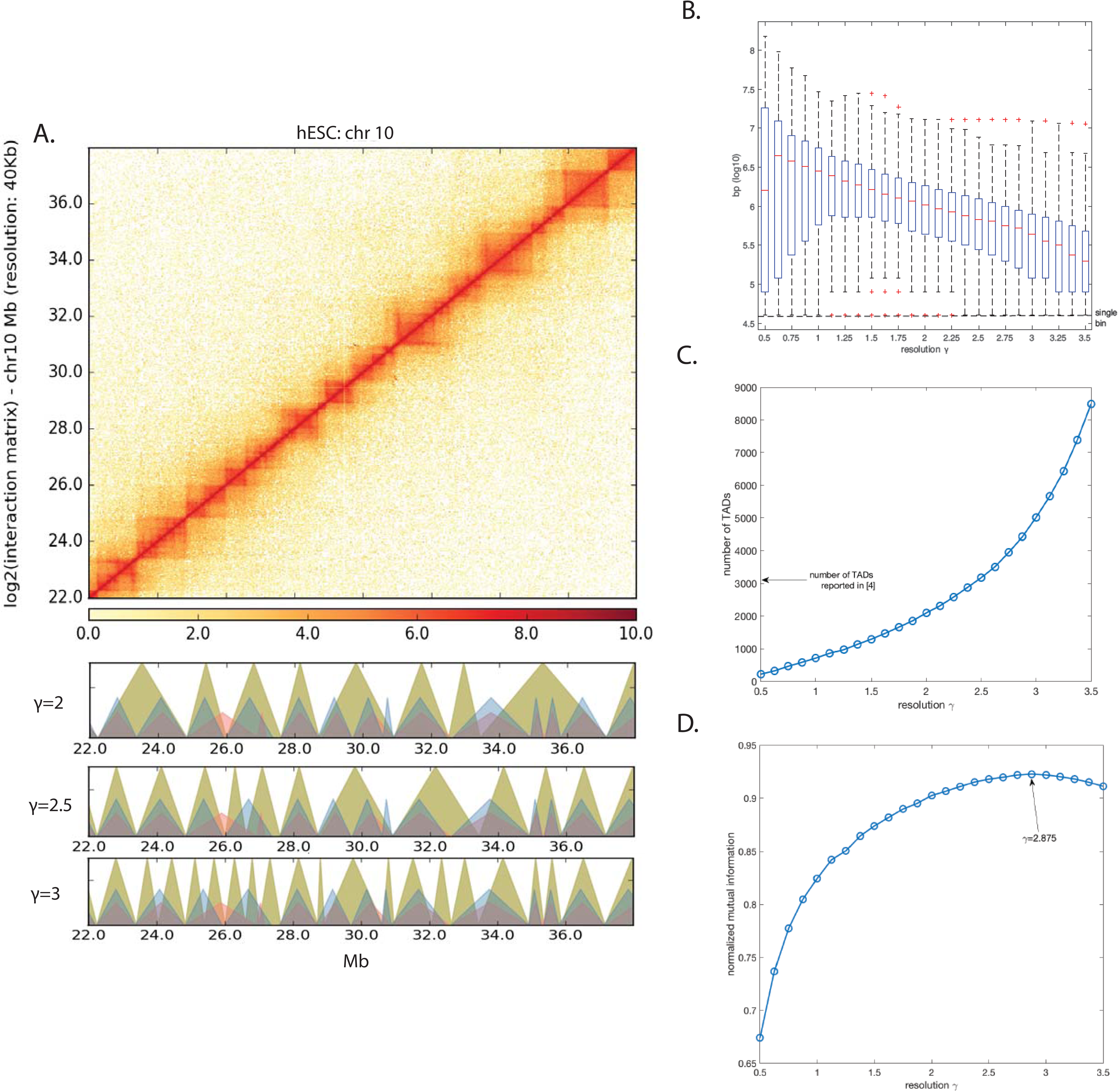
Identification of TADs in multiple resolutions. A) A part of the contact map of the chromosome 10 in hES cell. The greenish triangles below represent TADs called by MrTADFinder in three different resolutions. The TADs called agree well visually with the contact map. The blue triangles and red triangles represent TADs called in human ES cells and human IMR90 cells respectively as reported in [8]. B) The size of TADs called in different resolutions. The median TADs size decreases from 3 Mbp to 300 kbp as the resolution increases from 0.75 to 3.5. C) The number of TADs increases as the resolution increases. When *γ*=2.25, there are about 2600 TADs in hES cells with a median size of roughly 1Mb. The median size goes down to 300kb when the resolution increases to 3.5. The number of TADs identified in [8] is marked by the arrow. D) Comparing TADs called by MrTADFinder with TADs called in [8]. Two algorithms agree the most in a particular resolution (*γ* ≈ 2.875).

We then further compared the TADs identified at different resolutions by MrTADFinder with TADs identified by a previous method. As quantified by the normalized mutual information (see Methods for details), TADs identified by MrTADFinder best match with TADs identified in [8] when the resolution parameter is 2.9. In general, unless the resolution is sufficiently small (*γ* < 1.5), the two methods are quite consistent (see Figure 2D). Nevertheless, the introduction of the resolution parameter *γ* opens an extra dimension in domain identification in a sense the algorithm used in [8] focuses on a particular resolution instead.

### Signatures near TAD boundaries identified in various resolutions

The interplay between 3D genome organization and various chromatin features has widely been investigated since some of the first Hi-C experiments were reported [5][8][9]. Nevertheless, there is no clear-cut pattern emerges by aligning a variety of chromatin features with TADs (Figure S3), even though the occurrence of sharp peaks at the boundaries is quite apparent. By identifying TADs and their boundaries using MrTADFinder, we found the boundary signatures that are consistent with the observations previously reported [8], for instance, the enrichment of active promoter mark H3K4me3 or active enhancer mark H3K27ac, as well as the depletion of transcriptional repression mark like H3K9me3 (Figure 3A and Figure S4). To better understand the relationship between domains organization and different chromatin features, we further examined the chromatin features near different sets of boundaries that were identified in different resolutions. We found that in general, the enrichment of peak density at boundary decreases as resolution increases. This is because the number of TADs increases as the resolution increases, various chromatin features appear in the boundaries of low-resolution TADs do not appear in high-resolution TADs (Figure 3A). More specifically, the enrichment of histone marks like H3K36me3 and H3K4me3 exhibits a monotonic drop whereas certain marks exhibit characteristic resolutions. For instance, the enrichment of mark H3K27me3 remains high up to a resolution of *γ* = 2.5 (Figure 3B). The observation suggests that the mark H3K27me3 in general marks the boundary of TADs up to a particular resolution (length scale).

Beside epigenetic signatures, we examined the distribution of protein-coding genes along chromosomes in relation to TAD boundaries formation. Though the starting positions of genes tend to be enriched near TAD boundaries, the enrichment is much stronger for housekeeping genes as compared to tissue-specific genes (Figure 4A). As housekeeping genes are essentially active, the pattern resembles the active promoter mark H3K4me3 shown in Figure 3B. The discrepancy between housekeeping genes and tissue-specific genes was firstly reported in Ref. [8]. Nevertheless, by extending the idea to multiple resolutions, we found that the distribution of housekeeping genes follows a different length scale compared to tissue-specific genes. As shown in Figure 4B, housekeeping genes in general marks the boundary of TADs up to the resolution *γ* = 1.5.

**Figure 3.**
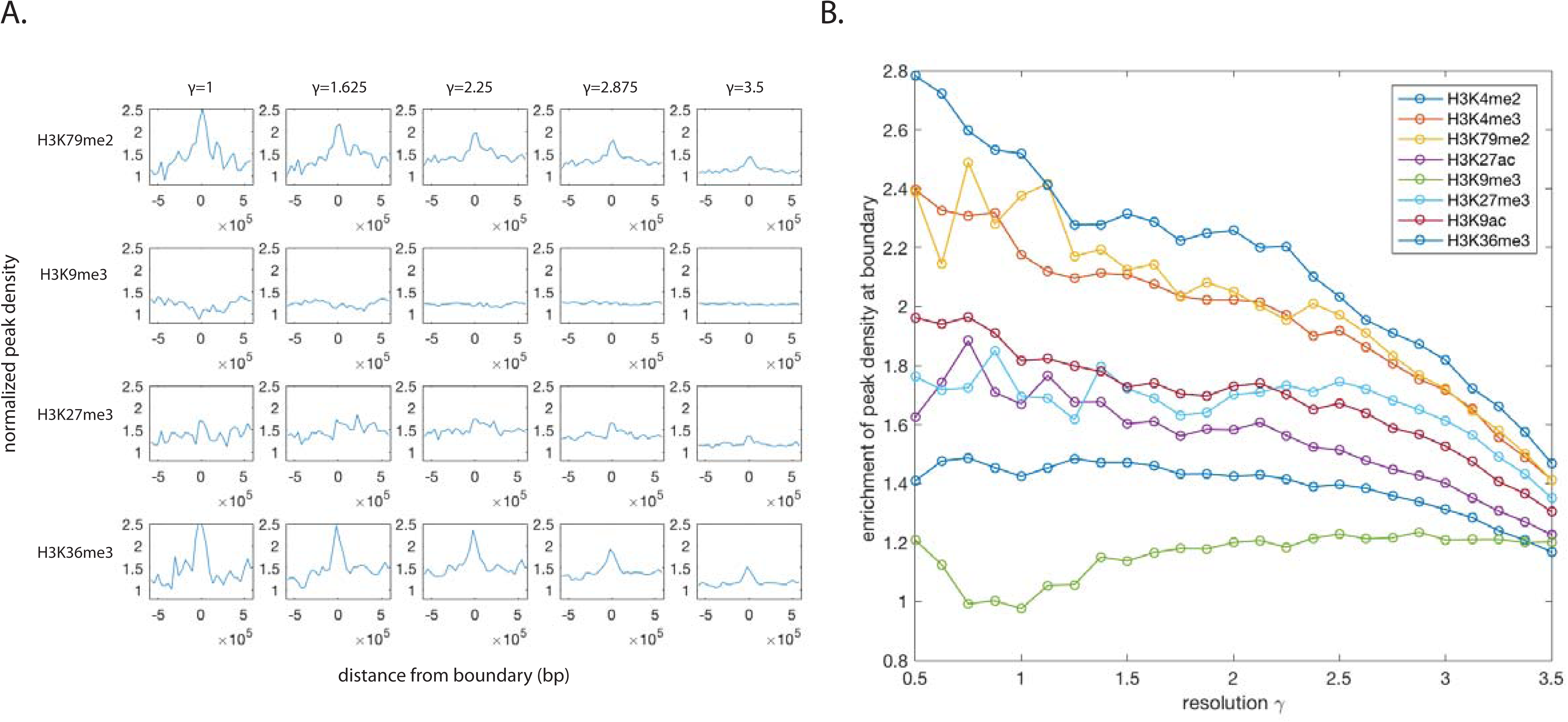
Boundary signatures of histone modifications in different resolutions. A) Histone modifications near the TAD boundary regions obtained in various resolutions. The peak density is obtained by counting the number of peaks in every 40kb bin, and normalized by a null model in which peaks are randomly distributed. B) Different histone marks show different levels of enrichment near TAD boundaries at different resolutions. Despite a general decreasing trend, the signal of certain marks likes H3K27me3 remains flat until a very high resolution.

**Figure 4.**
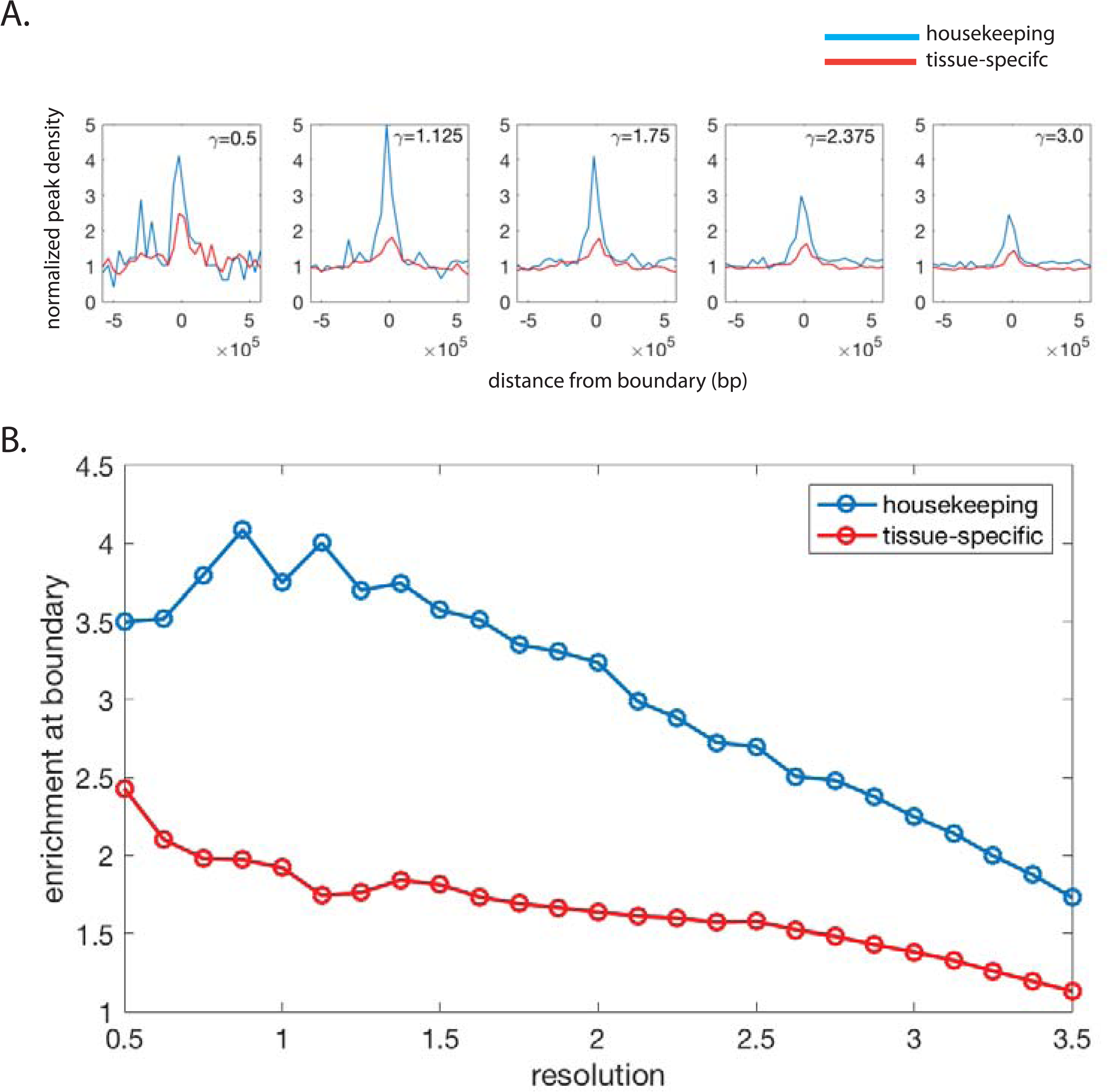
A) Distribution of house-keeping genes and tissue-specific genes near TAD boundaries at different resolutions. House-keeping genes are more enriched near TAD boundaries as compared to tissue-specific genes. B) House-keeping genes and tissue-specific genes show different levels of enrichment near TAD boundaries at different resolutions. Tissue-specific genes show a general decreasing trend, whereas the number of house-keeping genes remains flat until a high resolution.

### Binding of transcription factors near TAD boundaries identified in various resolutions

Apart from histone modifications, it is well known that certain transcription factor binding sites are enriched near the boundary regions of TADs [8]. Instead of looking at individual factors, we further explored the location of the so-called HOT regions and XOT regions on TADs. High-occupancy target (HOT) regions and extreme-occupancy target (XOT) regions are genomic regions that are bound by an extensive amount of transcription factors [16]. As expected, we found a strong enrichment of HOT regions and an even stronger enrichment of XOT regions near TAD boundaries in hES cells (Figure 5A). The observation is, in general, true for all tested resolutions. The observation agrees with the idea that HOT regions are very accessible regions in open chromatin. Nevertheless, it is still widely unknown if transcription factors bind to HOT regions simply because of thermodynamics, or the binding will result in important biological consequences.

**Figure 5.**
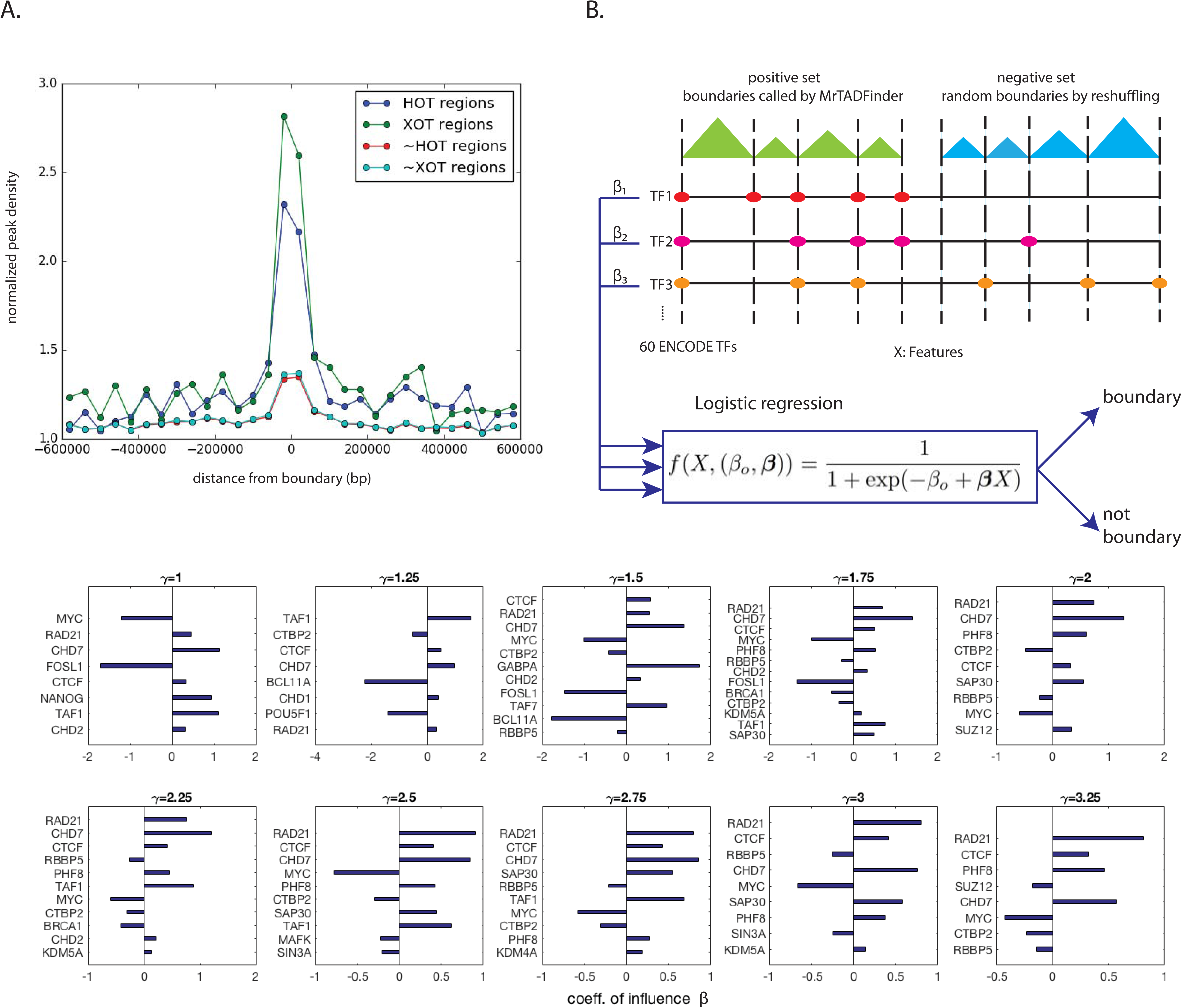
Transcription factors binding in different resolutions. A) Enrichment of HOT (high-occupancy target) and XOT (extreme-occupancy target) regions near TAD boundaries in hES cell. Boundaries are identified by MrTADFinder at a resolution *γ* = 2.75. The y-axis is normalized by a null model that peaks are randomly distributed in along the chromosome. B) A logistic regression model to classify real TAD boundaries and random boundaries based on thew binding pattern of 60 TFs. The most influential factors responsible for TAD boundaries formation at different resolutions are listed. Factors with a positive coefficient have a direct effect on border establishment or maintenance, whereas factors like MYC has a negative effect. The factors are sorted by corresponding P-values and only the significant factors are displayed.

Motivated by the observation that many factors tend to bind to the boundary regions, we further examine which factors are responsible for establishing the domain border, and more interestingly for borders in different resolutions. There are a few proteins which are widely known to be important in border establishment [17]; nevertheless, it is worthwhile to perform a systematic analysis. To do so, we formulated a classification problem which aims to distinguish, for each resolution, a set of boundaries identified by MrTADFinder (positive set) from a set of random boundaries obtained by swapping the TADs along the chromosomes (negative set). Using a logistic regression model recently proposed by [18], we integrated the binding signals of 60 transcription factors at a genomic locus to predict if it is TAD boundary (see Figure 5B and Methods for details). Generally speaking, with 10-fold cross validation, the model is quite successful in low resolutions (AUC=0.81, Figure S5). The result is consistent with an early work based on histone modifications [19]. Being consistent with the trend that chromatin features are less enriched at the boundaries of high resolution TADs, the predicting power of the model decreases as the resolution increases. The regression model further quantifies explicitly the influence of each of the transcription factors. In general, factors that are responsible for border formation are quite consistent across different resolutions (Figure 5B). For instance, we found that the well-known insulator CTCF, and Rad21 that is a part of cohesin, are direct key components of border establishment. In addition, the chromatin remodeler Chd7, which is often found at enhancers [20], is predicted to be a key component. On the other hand, factors like MYC have a consistently negative effect. Nevertheless, the relative importance of factors does change with resolutions. For instance, Rad21 has a higher predictive power in classifying high-resolution domains in compared with classifying low-resolution domains.

### Different resolutions suggest enhancer-promoter linkages in different length scales

The contact maps of more deeply sequenced Hi-C experiments have exhibited a pattern that a large fraction of TADs has “peaks” in their corner [21], meaning the contact frequency between the endpoints of such domains is higher than those of their surrounding neighborhood. The configuration suggests that the boundaries of such domains form a chromatin loop. We investigated if a similar conclusion could be drawn from the TADs called by MrTADFinder using a set of significant long-range promoter contacts identified by capture Hi-C [22]. Based on the Hi-C data of GM12878 in [21], we found that there are indeed potential promoter-enhancer linkages connecting the endpoints of domains. Moreover, by increasing the resolution parameters, the boundaries of the smaller TADs further capture the potential promoter-enhancer linkages in shorter length scales (Figure 6). It is worthwhile to point out that the linkages connecting the endpoints of domains form a small fraction as compared to the total number of significant interactions identified by capture Hi-C. Therefore, identifying the domain borders is not a direct method to predict potential enhancer-gene linkages. On the other hand, though the increase in the number of boundaries can capture a higher number of potential interactions, the same analysis for an ensemble of randomly reshuffled TADs shows the observation in TADs called by MrTADFinder is significant (Figure 6). In other words, TADs in a higher resolution are potential subTADs that mediate long-range interactions in a finer length scale [23].

**Figure 6.**
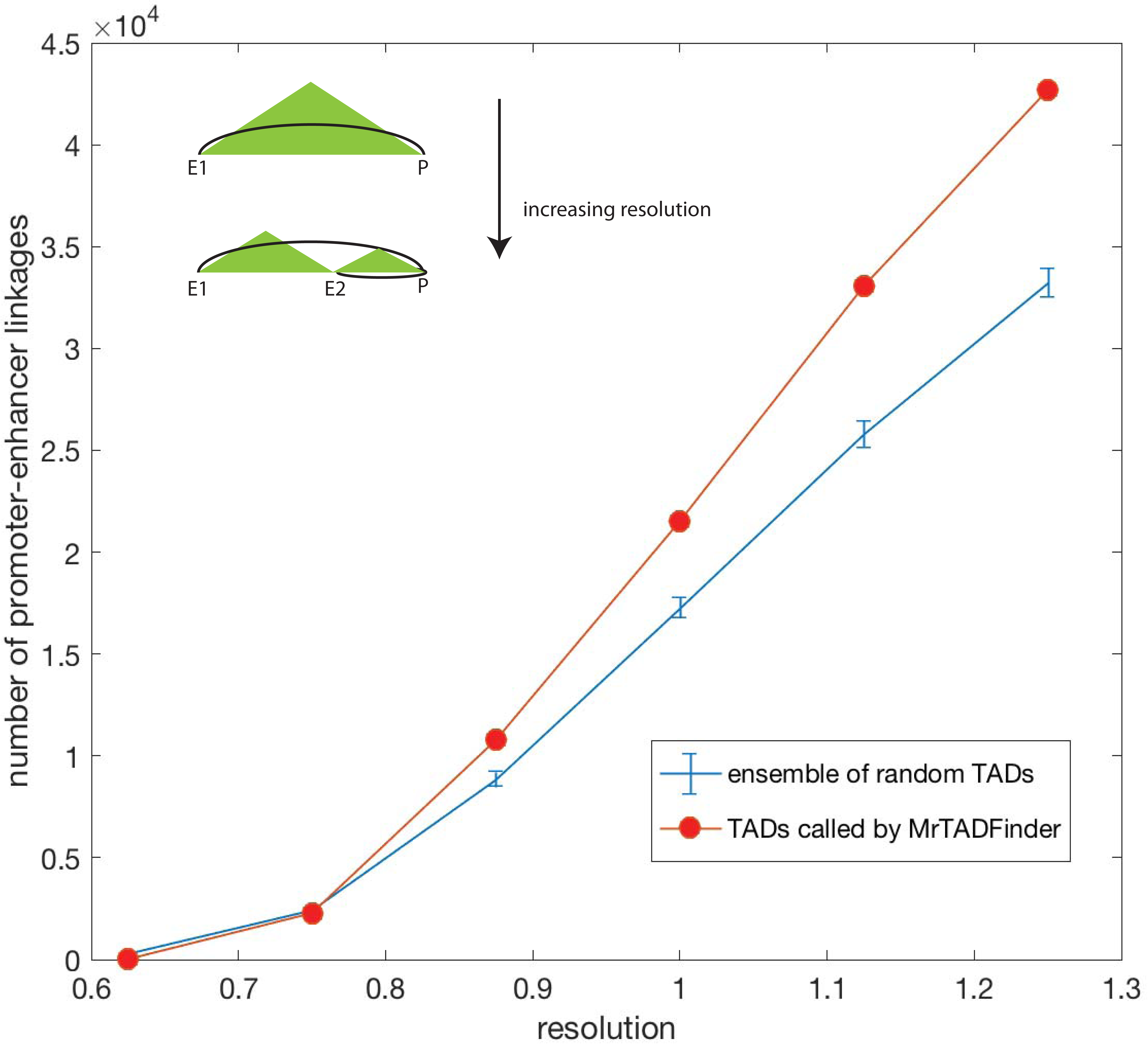
The number of promoter-enhancer linkages connecting the endpoints of domains in different resolutions. As the resolution increases, the increase in the number of boundaries can capture a higher number of potential interactions. The blue curve shows the increase for an ensemble of randomly reshuffled TADs. The number of promoter-enhancer linkages connecting the endpoints of real domains is higher than the random counterparts.

### TAD boundaries and mutational burden

We have examined the interplay between domains organization and chromatin features. Recently, it has been reported that epigenomic features shape the mutational landscape of cancer [24]. Motivated by this linkage, we further investigated the occurrence of somatic mutations near the boundaries. More specifically, we mapped the somatic mutations obtained from breast cancer samples to the TAD boundaries we identified in MCF7 cells (see Methods). In a given resolution, there are 85 boundary regions identified on chromosome 10. The regions can be clustered into 3 groups based on the positional distribution of somatic mutations. As in shown in Figure 6, two of the clusters exhibit a step-function behavior (blue and red) in which the abrupt transition essentially happens at the boundary. For boundary regions in the remaining cluster, the mutational burden exhibits no difference across the TAD boundaries. Because of the close relationship between TADs and replication-timing domains [25], the observation resonates with a well-known observation that genomic regions with a high mutational burden are replicated at a later stage during DNA-replication [26]. As shown in the inset, using Repli-seq data in S1 phase, the upstream regions of the boundaries found in the blue cluster have a high mutation rate but a low Repli-seq signal, meaning they are indeed replicated at a later stage during replication. On the contrary, the upstream regions of the boundaries found in the red cluster are replicated at an early stage and therefore exhibit a low mutation rate.

Motivated by the relationship between TADs and DNA replication, we overlaid TADs in different resolutions with data from Repli-seq experiment (Figure S6). We observed that TADs identified in different resolutions match with the Repli-seq data in different stages of a cell cycle. For instance, while a TAD identified in a low resolution does not replicate at an early phase, say S1, its sub-structures identified in a higher resolution correspond to two separate peaks at later stages, say S2 and S3 (Figure S7). Nevertheless, it is worthwhile to point out that mapping Hi-C reads from cancer cell lines like MCF7 to the reference genome is not perfect because quite some reads may come from translocations or copy number variations. Computational approaches have recently been developed to perform correction as well as to infer those large scale genomic alterations [27][28].

### Comparison with existing methods based on CTCF enrichment

There are quite a few existing methods on identifying TADs using Hi-C data. Dixon *et al*. identified TADs based on the so-called directionality index using Hi-C data in hES cell and found an enrichment of CTCF binding sites at the boundary regions [8]. Since then the enrichment of chromatin features has been used as a benchmark for various TAD calling algorithms [29][30][31]. As a comparison, we performed the same analysis using TADs based on MrTADFinder. As shown in Figure 7, both methods exhibit a similar pattern. In fact, as reported in [29][30][31], the enrichment pattern of CTCF binding peaks is qualitatively the same for all the proposed methods. By repeating the analysis in different resolutions, we observed that the level of enrichment depends on the resolution (Figure 7, Figure S7). At a low resolution, i.e. for larger TADs, the enrichment signal is stronger, and the signal tends to extend over a longer distance from the boundary. At a higher resolution, the signal is weaker and confined to near the boundary. In general, Figure 7 suggests that boundaries identified in lower resolutions are more likely to be bound by CTCFs. From a biological standpoint, as a boundary identified in a lower resolution separates two large domains, the results may bring insights on how to mediate chromatin loops at different length scales via an important architectural protein [32][33]. As the level of CTCF enrichment might be the consequence of different chromatin length scales, it might not be fair to use it directly for benchmarking the performance of different algorithms.

**Figure 7.**
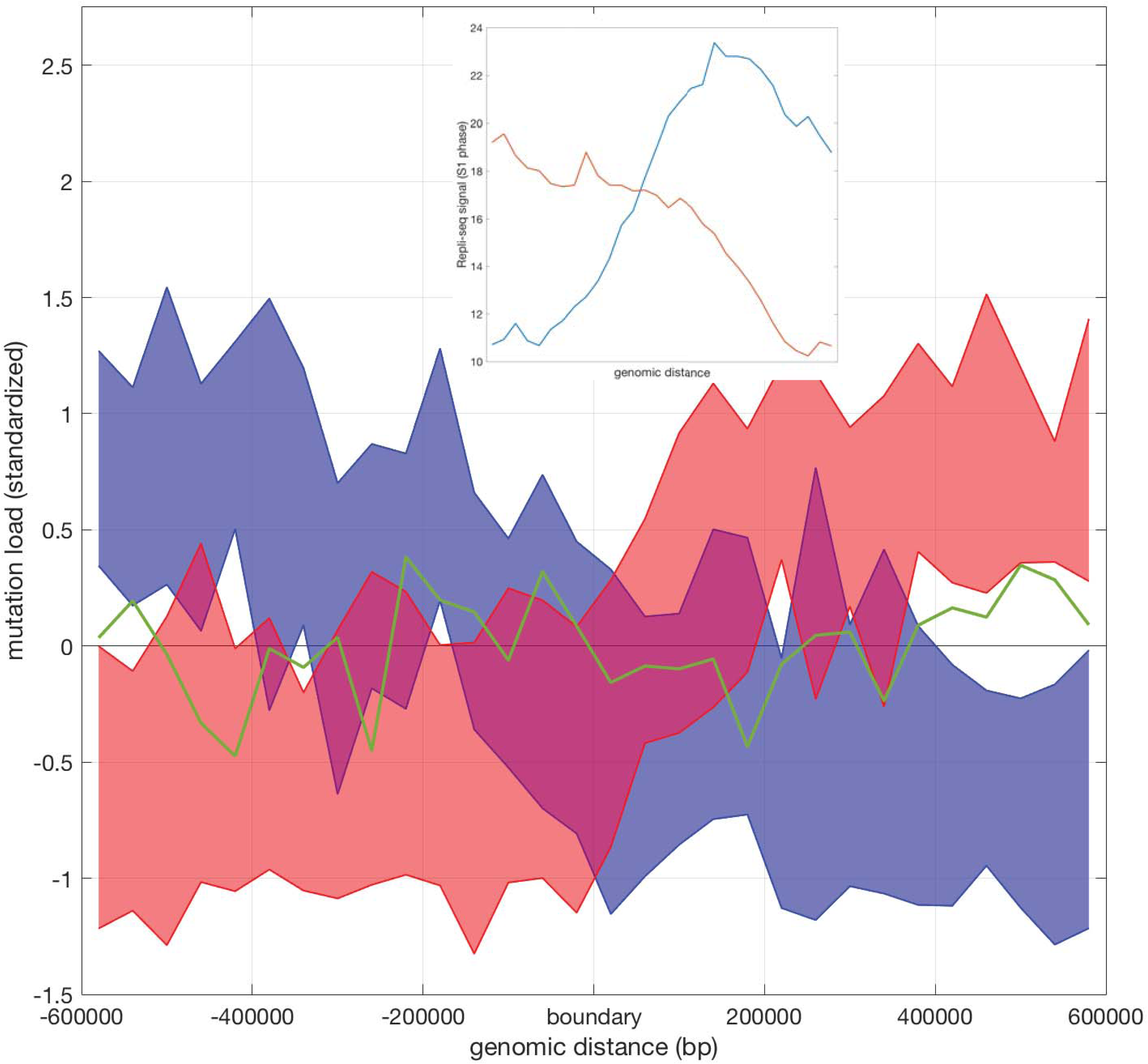
Mutational burdens across TAD boundaries. The 3 clusters of boundary regions exhibit distinct patterns in terms of mutational burden. For blue and red clusters, the area marks the first and the third quartiles. For the green cluster, only the mean values at different positions are shown for clarity. The inset shows the average Repli-seq signal for the red and blue clusters.

### Robustness, performance and implementation of MrTADFinder

Because of the stochastic nature of the modified Louvain algorithm, we explored the robustness of MrTADFinder. In the current setting based on multiple runs of the modified Louvain procedure, we found the results of two independent callings highly robust. In fact, the normalized mutual information is 0.99 (see Figure S8). We further investigated how MrTADFinder performs in replicates. Using Hi-C data released by the ENCODE consortium, we found that TADs called in a pair of biological replicates agree reasonably well, with normalized mutual information about 0.85 (see Figure S9 and Methods).

MrTADFinder is implemented in Julia. Julia programmers can import MrTADFinder as a library for calling various functions. It can also be run in command line if Julia and the required packages are installed. The performance of MrTADFinder, in general, depends on the size of the input contact map. We have tested the performance using the contact maps of GM12878 cell generated by the Aiden lab [21]. The performance is reasonable. For instance, for chromosome 10, in a bin-size of 25kb (i.e. a contact map 5400 by 5400), the time required to arrive at all TADs with 10 runs of Louvain algorithm is about 20 minutes on a laptop with 2.8GHz Intel Core i7 and 16Gb of RAM. The time required is only 6 minutes if the bin size is 50kb. We have made the source code available on GitHub (see software availability).

### Optimization based on recurrence relation

Despite the similarity between equations (1) and (2), network modules are rather arbitrary collections of nodes, but domains are continuous segments along the chromosome. In fact, the total number of possible partitions for a chromosome is much smaller than the total number of ways to divide a network into modules. As a result, while the optimization of equation (1) is an NP-hard problem, the optimization of (2) can be quite efficiently solved using a dynamic programming inspired method (see Methods and Figure S10). It is instructive to explore this avenue because quite some algorithms for identifying TADs are based on a similar approach but with different objective functions [29][30][31]. Moreover, by enumerating all possible ways to decompose a chromosome into TADs, one could write down the partition function and define a probability of occurrence for each of the possible partition in a statistical mechanics’ manner.

The time complexity of this algorithm is in order of ?(*n*^3^), where n is the size of the contact map. Given the time complexity, finding the optimal partition using a bin size of 40kb is quite impractical. For instance, the calculation takes about an hour for chromosome 21, as compared to seconds by using the heuristic. Therefore, though the connection between identifying TADs and problems like finding RNA secondary structure is of theoretical interest, MrTADFinder is developed based on the modified Louvain algorithm. Nevertheless, we have implemented the approach based on recurrence relation and performed a comparison with the heuristic. Using a contact map of hES cell (chromosome 1) with a bin size of 500kb, we found the sub-optimal partitions based on our modified Louvain algorithm are very close to the optimal partition. The normalized mutual information between optimal and sub-optimal values is 0.977±0.007.

## Discussion

In this paper, we have introduced an algorithm to identify TADs from Hi-C data and performed several analyses to show the biological significance of the TADs identified. In particular, by introducing a single continuous parameter *γ*, we can further examine domains organization and its interplay with a variety of chromatin features in multiple resolutions. It is important to emphasize that the idea of resolution we introduced in MrTADFinder is different from some other usages of the same term in Hi-C analysis. From an experimental standpoint, the resolution of a Hi-C experiment refers to the average fragment size as digested by restriction enzymes (∼4kb to ∼1kb) [5][21] or more recently by micrococcal nuclease (∼150bp) [34]. Regarding the construction of contact maps, the term resolution has been used to refer to the bin size, where the proper choice usually depends on the number of reads in the stage of data processing. Both usages are primarily technical. What we mean by resolution, however, refers to the multiple length scales built inside the organization of the genome. It is well known that there are structures in different length scales such as compartment, domains, and sub-domains [35], and chromatin features like histone marks exhibit multiple length scales [36]. The concept of resolution introduced here points to the integration of these structures and enables one to explore the rich structures hidden in contact maps. From a practical point of view, *γ* = 1 seems to be the natural starting point. One could increase or decrease the value of *γ* in order to explore the intrinsic structure. Nevertheless, because of the different contact maps might have various differences like the read coverage, one should be cautious to directly compare the resolution parameters between different contact maps.

A novel contribution of this work is the derivation of an expected model for any intra-chromosomal contact map by solving a system of matrix equations. The null model preserves the coverage of each genomic bin as well as the distance dependence of contact frequencies in the observed map. As such features of contact maps are involved in most computational analysis of Hi-C data, apart from the identification of TADs, the expected model can be used for applications like finding compartments [5] and identifying potential enhancer-target linkages [37]. Mathematically, the expected matrix is solved by an iterative procedure. The procedure can be regarded as a generalization of a class of matrix balancing methods used for normalizing Hi-C matrices [38], as the later is merely a different set of matrix equations. However, it is important to emphasize that the so-called ICE algorithm aims to remove bias in the contact map, whereas our method aims to generate a background model. While MrTADFinder focuses on intra-chromosomal interactions, recent studies employ various clustering methods to identify inter-chromosomal clusters using Hi-C contact frequency [39][40]. It is worthwhile to point out that similar expected models used in this study can also be derived for inter-chromosomal interactions to better separate signal and noise.

Several methods have been developed for identifying TADs from Hi-C data [41]. One of the earliest methods is based on the so-called directionality index, a 1D statistic measuring whether the contacts have an upstream or downstream bias [8], and later the bias is exploited by the so-called arrowhead algorithm [21]. Later algorithms exploit the block diagonal nature of TADs in a contact map [29] [30][42]. Though some of these algorithms do take the distance dependence into the background, but they do not take into account both the genomic distance and the effects of coverage in a compact mathematical formalism. The algorithm TADtree [30], and more recent efforts, namely Matryoshka [31] and metaTAD [43] aim to investigate the hierarchical organization of TADs based on a tree structure. Indeed, merging smaller TADs at the lower level of the hierarchy results at larger TADs similar to the TADs obtained by MrTADFinder at a low resolution. Nevertheless, MrTADFinder does not impose a hierarchical organization. The probabilistic nature of Louvain algorithm enables the definition of TAD boundaries in a probabilistic fashion, and therefore a possibility to define overlapping TADs. To a certain extent, the idea of continuous resolution used in MrTADFinder is distinct in comparison with algorithms based on a bottom-up approach, but similar in spirit to Ref. [29].

MrTADFinder is motivated by the community detection problem in network studies. Although a network perspective of chromosomal interactions has previously been proposed [44][45], a lot of widely studied concepts in networks have rarely been explored in the context of chromosomal organization. A network representation is arguably more flexible than a simple matrix representation, for instance, transcription factors binding and histone modifications can be easily incorporated into the network, forming a decorated network. Moreover, one could extend the framework by concatenating multiple Hi-C contact maps to form a multi-layer network. The same idea has been used for cross-species transcriptomic analysis [46]. By facilitating the application of a variety of graph-theoretical tools, we believe that network algorithms will be useful for future studies on the spatial organization of the genome.

## Materials and methods

### Hi-C data and their pre-processing

The Hi-C data of human ES cells and IMR90 cells were reported in Ref. [8]. Raw reads were processed using Hi-C Pro [47], arriving at contact matrices in various bin sizes. In all analysis, the whole-genome contact map was iteratively corrected for uniform coverage [38]. Intra-chromosomal contact maps were then extracted from the whole-genome contact map of bin size 40kb for downstream analysis. Hi-C data and contact maps in MCF7 cells were reported in Ref. [48]. The whole-genome contact map provided was binned with 40kb bin size and was already passed the ICE normalization. Data in GM12878 were reported in [21], with bin size 25kb. The ENCODE Hi-C data were released by the ENCODE consortium. Altogether 8 cell lines with a relatively higher coverage were used in the reproducibility analysis including T47D, A549, Caki2, G401, NCI-H460, Panc1, RPMI-7951 and SK-MEL-5. For each cell line, two replicates were separately used. The ENCODE Hi-C data were processed by the tool cworld (https://github.com/dekkerlab/cworld-dekker). Capture Hi-C data were reported in Ref. [22]. Only 1618000 significant interactions linking promoters and non-promoter regions were included in the analysis of Figure 8. Visualization of contact maps were all generated by the tool HiCPlotter [49].

**Figure 8.**
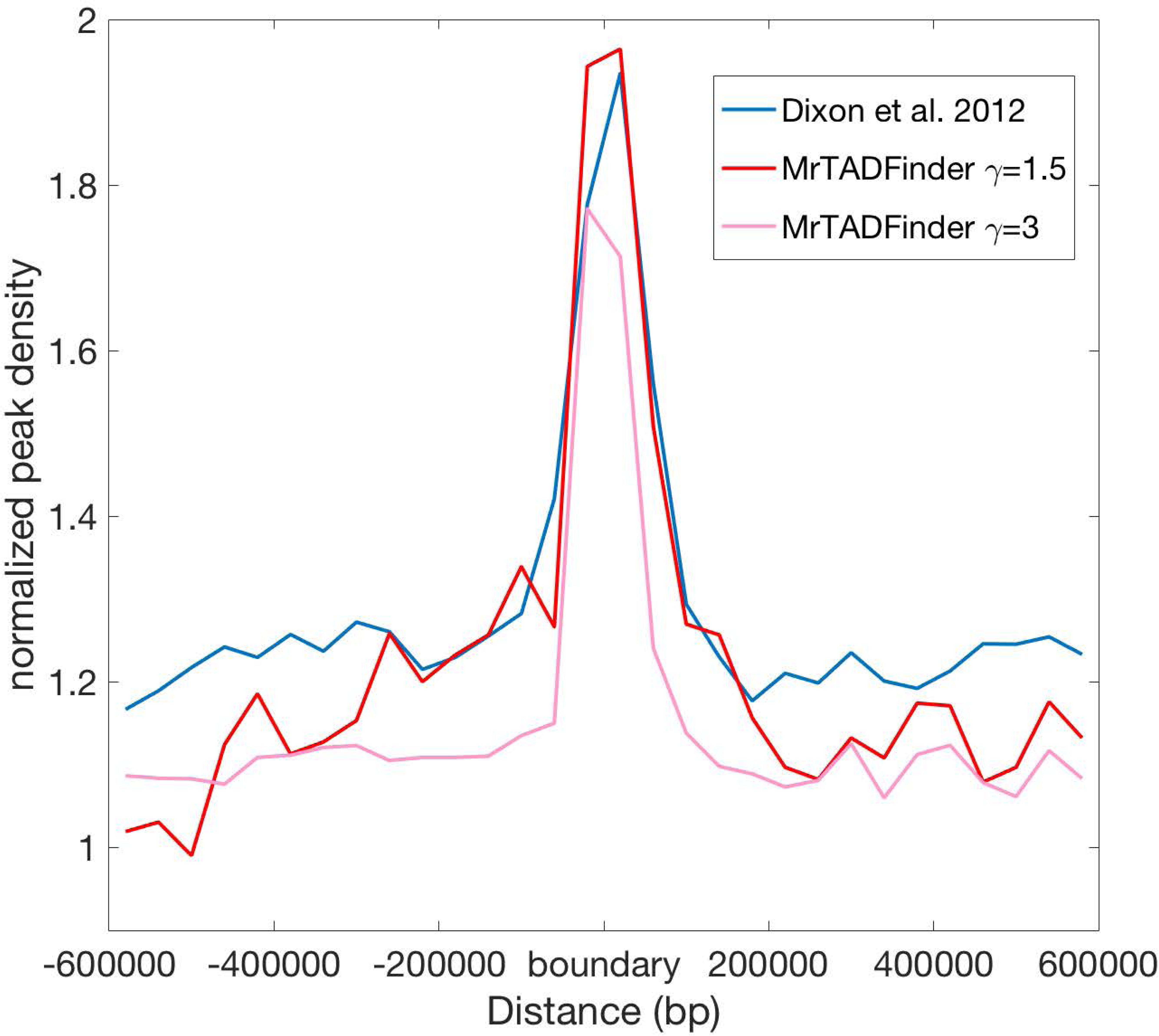
Enrichment of CTCF peaks near TAD boundaries at two different resolutions. The red line shows the same analysis using TADs reported in [8].

### Chromatin Data

All chromatin data, including histone modifications, transcription factors binding, expression, replication timing, were downloaded from the ENCODE data portal.

### Deriving a background model for any given intra-chromosomal contact map

The average number of contacts as a function of genomic distance can be estimated by considering all elements in matrix *W.* A local smoothing approach similar to the method used in [50] was employed. The window size equals to 1% of the data.

Equation (3) and (4) can be rewritten in the form

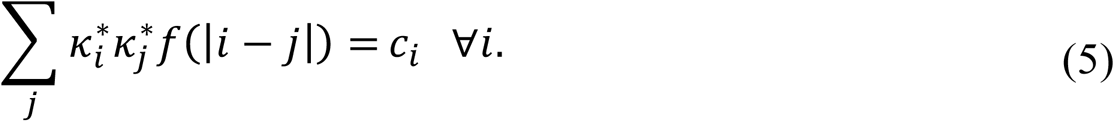

The system of non-linear equation is similar to the matrix balance approach used in [38]. As the aim of [38] is to remove bias, the coverage *c*_*i*_ is the same for all bin *i* and *f* is replaced by the original empirical map. Nevertheless, the unknowns 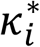 can be used by a similar iterative procedure as proposed in [38].

### Heuristic procedures for optimizing Q

To optimize the objective function *Q*, we employ a modified version of Louvain algorithm [15], which is widely used in identifying modules in networks (see Figure 1). In a nutshell, the algorithm consists of two steps. The algorithm starts as every bin has its own label, and the label will end up as an identifier for the module it belongs. In the first step, for each bin, we update its label by either choosing the label of one of its two neighboring bins or by remaining unchanged based on whether or not the value of *Q* will be increased. There will be multiple rounds of updates in this step. For each round of update, we go through all the bins once, but the order is random. The updating procedure will be repeated for multiple rounds until no more update is possible. We will then perform the second step such that the bins with the same labels will be locked together, in a sense their labels will only be updated in a synchronized fashion. It is worthwhile to mention that the updating procedure in the first step makes sure bins with the same labels form a continuous segment. Once the bins are locked to form super-bins, the first step will be performed again but in the level of super-bins. The two steps will be repeated iteratively until no increase of modularity is possible.

The output of the modified Louvain algorithm is essentially a particular partition of the entire chromosome. As the result of the algorithm, in general, depends on the order of updates, multiple runs are performed to probe the fuzziness of the assignment. As the chromosome is binned into *n* equally sized bins, we examine, say after 10 trials, how likely the border between bin *i* and bin *i* + 1 is indeed a domain boundary, i.e. bin *i* and bin *i* + 1 are called to belong to two different TADs by the modified Louvain algorithm. We then naturally define a boundary score for each of the *n*+1 borders as the fraction of trials in which a border is called as a boundary. To define a set of consensus boundaries, we choose a cut-off of 0.9. In other words, the border between two adjacent bins is defined as a confident boundary only if they are called to belong to two different domains in at least 9 out of 10 trials. The final output of MrTADFinder is a set of consensus TADs defined as regions between the consensus domains

The boundary score assigned to each border is not merely an immediate but serves as a proxy of the degree of insulation. A border with a high boundary score is more effective in forbidding the contacts between its left and right regions.

### Quantifying the consistency between two sets of TADs

Given two sets of TADs, say in different cell lines, or called by different algorithms, we employ the so-called normalized mutual information to quantify the consistency. Suppose *X* and *Y* are two random variables whose values *x*_*i*_ and *y*_*i*_ represent the corresponding domain labels of bin *i*. The normalized mutual information MI_norm_ is defined as

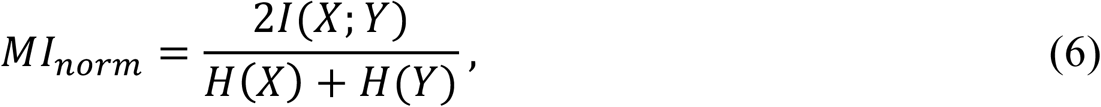

here *H*(*X*), *H*(*Y*) are the entropy of *X* and *Y*, and *I*(*X*; *Y*) is the mutual information quantifying to what extent the domain labels in *X* predict the labels in Y. A normalized form of mutual information is used here to make sure the value lies between 0 and 1 for comparison. To have a fair comparison, bins that are not assigned to any TADs in both sets of partitions are not counted. If two sets of partitions are identical, the value of normalized mutual information is 1.

### Chromatin signatures within TADs in different resolutions

Given the location of binding peaks of a transcription factor or a histone mark, the peak density near TAD boundaries was estimated by considering for all boundaries the region from upstream 600kb to downstream 600kb. The regions were aligned, and the number of peaks was summed accordingly. To calculate the enrichment, the number of peaks was normalized by the expected number of peaks in a particular region under a null model that peaks are randomly distributed in the genome.

The influence of individual transcription factors on the formation of domain borders was formulated as a classification problem. For a particular resolution, the set of boundaries called by MrTADFinder was used as a positive set whereas a set of random boundaries obtained by swapping the TADs along the genome was chosen as the negative set. The signal values of 60 transcription factors are used as features for classification. The combined effect of all features was modeled the logistic function

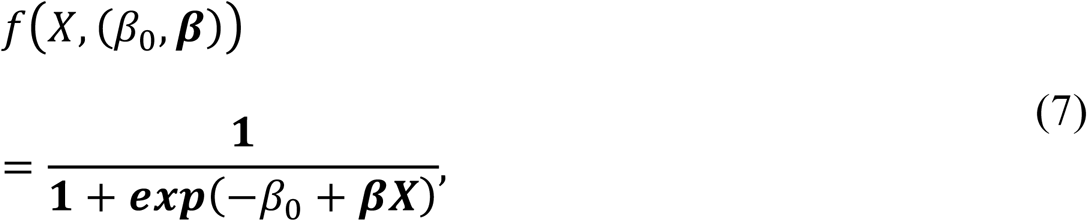

here X represents all features; *β* is a vector determining the coefficients of influence for all features and *β*_*o*_ is a bias parameter. Given a training set, a likelihood function was defined. An optimal *β* was inferred by optimizing the likelihood function using gradient descent with L1-regularization. The inferred logistic function was used to predict the test set. To have a more accurate estimate, 10-fold cross-validation was performed, and the error bars were estimated by multiple negative training sets.

### Somatic mutations

The set of somatic mutations were downloaded from the data portal of the International Cancer Genome Consortium (ICGC). The mutations were called the breast cancer samples of 676 donors. The samples were sequenced in a whole-genome level. Breast cancer samples were used in this analysis to match the Hi-C data of MCF7 cell.

### Optimal partition

The idea is to extensively enumerate all the possible partitions of the chromosome. In a nutshell, a binned chromosome can be considered as a sequence (1, 2,…, *n* - 1, *n*). Rather than partitioning the whole sequence at a first place, we look for the optimal partition for all the possible sub-sequences starting from sub-sequences with length 1. Let us denote the optimal value of modularity *Q* for a sequence *a*_1_*a*_2_ … *a*_*l*–1_*a*_*l*_ as *optQ*(*a*_1_*a*_2_*… a*_*l*–1_*a*_*l*_). The value is the maximum of the following *l* possibilities:

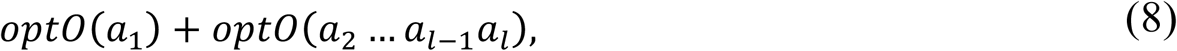

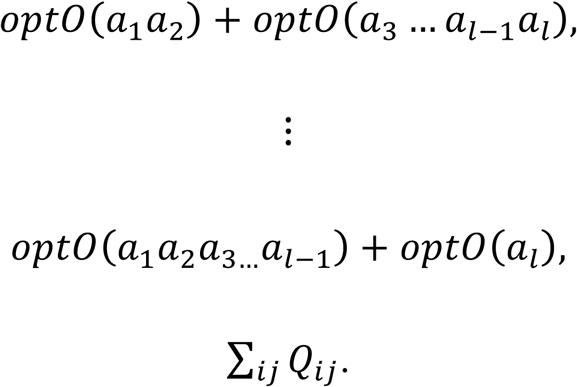

Suppose the maximum is the sum *optO*(*a*_1_*a*_2_ … *a*_*r*_) + *optO*(*a*_*r*+1_ … *a*_*l*–1_*a*_*l*_), where 1 ≤ *r* < *l*. The sum corresponds to the case that the optimal partition of *a*_1_*a*_2_ … *a*_*l*_ is a combination of the optimal partitions of *a*_1_*a*_2_ … *a*_*l*_ and *a*_r+1_ … *a*_*l*–1_*a*_*l*_ (see Figure S10). It is not necessary that *a*_1_*a*_2_ … *a*_*r*_ forms a single domain. The key is that the expression *optQ*(*a*_1_*a*_2_ … *a*_*l*–1_*a*_*l*_) can be found recursively because all possibilities depend on the optimal values of sub-sequences shorter than *l*. The last summation in (4) sums Q over all positions from *a*_1_ to *a*_b_, meaning the *l* bins belong to the same domain. Once the value of *optQ*(*a*_1_*a*_2_ … *a*_*n*–1_*a*_*n*_) is found, we can trace back the actual partition for the whole chromosome. As shown in the source code, it takes three loops to enumerate all possible partitions. The procedure is analogous to the Nussinov algorithm in finding the optimal secondary structure of RNA [51].

## Acknowledgments

We want to thank the 3D Nucleome subgroup in the ENCODE consortium for discussion and data processing. KKY acknowledges Anurag Sethi, Joel Rozowsky, Sushant Kumar and Arif Harmanci for feedback and discussion. KKY acknowledges Timur Galeev and Jonathan Warrell for critical reading on an earlier version of the manuscript. This work was supported by the HPC facilities operated by, and the staff of, the Yale Center for Research Computing.

## Author Contributions

Conceived and designed the study: KKY, with input from MG. Performed the research: KKY, SKL. Wrote the paper: KKY, MG.

## Software availability

The source code can be downloaded at https://github.com/gersteinlab/MrTADFinder.

## Supporting Information

Figure S1: Dependence of contact frequency and genomic distance. The analysis was performed using the contact map of the chromosome 1 of MCF7, binned in 250kb sized bins. The red line *f*(*d*) is the average contact frequency as a function of distance *d* obtained by smoothing. The green line shows a power-law function *d*^−1^.

Figure S2. Effective coverage 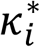 of loci is highly correlated with the coverage *c*_*i*_.

Figure S3. Aligning chromatin features with TADs in different resolutions.

Figure S4. Boundary signatures of 8 histone modifications in different resolutions (an extension of Figure 3A.)

Figure S5. Using transcription factors binding signals for predicting TAD boundaries. For each resolution, a logistic regression model based on transcription factors binding signals was trained to classify the TAD boundaries versus a set of random boundaries. The error bars were estimated by repeating the analysis using an ensemble of random boundaries. The performance (AUC and ACC) decreases as the resolution increases.

Figure S6. The relationship between TADs and DNA replication timing. TADs are identified for IMR90 using different resolutions. Signals of Repli-seq data in various stages of a cell cycle and a part of the contact map of the chromosome 10 are displayed. The TADs match visually well with the replication timing signals. The middle TAD identified in *γ* = 1 does not replicate at S1, its sub-units identified in *γ* = 1.25 replicate in S2 and S3.as shown by the peaks in the Repli-seq signal.

Figure S7. Enrichment of CTCF peaks near TAD boundaries at two different resolutions. The red line shows the same analysis using TADs reported in [8]. This figure is an extension of Figure 5.

Figure S8. Robustness of MrTADFinder. Histogram for pairs of independently called TADs. Using the default parameters (10 trials of the modified Louvain algorithm and a cut-off of 0.9), the normalized mutual information between two sets of called domains agrees extremely well (nMI=0.99).

Figure S9: Comparing TADs in biological replicates. For each cell line, TADs were called separately in each replicate for all chromosomes. The boxplot shows the distribution of the normalized mutual information for 23 chromosomes in different cell lines.

Figure S10: Identifying TADs by dynamic programming. The optimal value of Q for a chromosome segment running from ***i*** to ***j*** is stored in ***M***_***ij***_. The values of all elements in ***M*** can be enumerated using dynamic programming, starting from fragments of length 1 where ***M***_***ii***_ = ***Q***_***ii***_. There are different ways to divide a fragment of length l (gray lines). Suppose the optimal way is marked by the red line, then ***M***_1*l*_ = ***M***_1*r*_ + ***M***_*rl*_.

## References

[1] J. Dekker, M. A. Marti-Renom, and L. A. Mirny, “Exploring the three-dimensional organization of genomes: interpreting chromatin interaction data,” Nat. Rev. Genet., vol. 14, no. 6, pp. 390–403, Jun. 2013.

[2] V. I. Risca and W. J. Greenleaf, “Unraveling the 3D genome: genomics tools for multiscale exploration,” Trends Genet., vol. 31, no. 7, pp. 357–372, Jul. 2015.

[3] M. J. Rowley and V. G. Corces, “The three-dimensional genome: principles and roles of long-distance interactions,” Curr. Opin. Cell Biol., vol. 40, pp. 8–14, Jun. 2016.

[4] B. Bonev and G. Cavalli, “Organization and function of the 3D genome,” Nat. Rev. Genet., vol. 17, no. 11, pp. 661–678, Nov. 2016.

[5] E. Lieberman-Aiden et al., “Comprehensive Mapping of Long-Range Interactions Reveals Folding Principles of the Human Genome,” Science, vol. 326, no. 5950, pp. 289–293, Oct. 2009.

[6] R. Kalhor, H. Tjong, N. Jayathilaka, F. Alber, and L. Chen, “Genome architectures revealed by tethered chromosome conformation capture and population-based modeling,” Nat. Biotechnol., vol. 30, no. 1, pp. 90–98, Dec. 2011.

[7] M. J. Fullwood and Y. Ruan, “ChIP-based methods for the identification of long-range chromatin interactions,” J. Cell. Biochem., vol. 107, no. 1, pp. 30–39, May 2009.

[8] J. R. Dixon et al., “Topological domains in mammalian genomes identified by analysis of chromatin interactions,” Nature, vol. 485, no. 7398, pp. 376–380, May 2012.

[9] T. Sexton et al., “Three-Dimensional Folding and Functional Organization Principles of the Drosophila Genome,” Cell, vol. 148, no. 3, pp. 458–472, Feb. 2012.

[10] J. Dekker and E. Heard, “Structural and functional diversity of Topologically Associating Domains,” FEBS Lett., vol. 589, no. 20, Part A, pp. 2877–2884, Oct. 2015.

[11] A.-L. Valton and J. Dekker, “TAD disruption as oncogenic driver,” Curr. Opin. Genet. Dev., vol. 36, pp. 34–40, Feb. 2016.

[12] D. G. LupiÁñez, M. Spielmann, and S. Mundlos, “Breaking TADs: How Alterations of Chromatin Domains Result in Disease,” Trends Genet., vol. 32, no. 4, pp. 225–237, Apr. 2016.

[13] M. E. J. Newman, “Modularity and Community Structure in Networks,” Proc. Natl. Acad. Sci., vol. 103, no. 23, pp. 8577–8582, Jun. 2006.

[14] S. Fortunato and M. Barthélemy, “Resolution limit in community detection,” Proc. Natl. Acad. Sci., vol. 104, no. 1, pp. 36–41, Jan. 2007.

[15] V. D. Blondel, J.-L. Guillaume, R. Lambiotte, and E. Lefebvre, “Fast unfolding of communities in large networks,” J. Stat. Mech. Theory Exp., vol. 2008, no. 10, p. P10008, Oct. 2008.

[16] A. P. Boyle et al., “Comparative analysis of regulatory information and circuits across distant species,” Nature, vol. 512, no. 7515, pp. 453–456, Aug. 2014.

[17] E. Gómez-Díaz and V. G. Corces, “Architectural proteins: regulators of 3D genome organization in cell fate,” Trends Cell Biol., vol. 24, no. 11, pp. 703–711, Nov. 2014.

[18] R. Mourad and O. Cuvier, “Computational Identification of Genomic Features That Influence 3D Chromatin Domain Formation,” PLOS Comput Biol, vol. 12, no. 5, p. e1004908, May 2016.

[19] J. Huang, E. Marco, L. Pinello, and G.-C. Yuan, “Predicting chromatin organization using histone marks,” Genome Biol., vol. 16, no. 1, p. 162, Aug. 2015.

[20] M. P. Schnetz et al., “CHD7 Targets Active Gene Enhancer Elements to Modulate ES Cell-Specific Gene Expression,” PLOS Genet., vol. 6, no. 7, p. e1001023, Jul. 2010.

[21] S. S. P. Rao et al., “A 3D Map of the Human Genome at Kilobase Resolution Reveals Principles of Chromatin Looping,” Cell, vol. 159, no. 7, pp. 1665–1680, Dec. 2014.

[22] B. Mifsud et al., “Mapping long-range promoter contacts in human cells with high-resolution capture Hi-C,” Nat. Genet., vol. 47, no. 6, pp. 598–606, Jun. 2015.

[23] J. E. Phillips-Cremins et al., “Architectural Protein Subclasses Shape 3D Organization of Genomes during Lineage Commitment,” Cell, vol. 153, no. 6, pp. 1281–1295, Jun. 2013.

[24] P. Polak et al., “Cell-of-origin chromatin organization shapes the mutational landscape of cancer,” Nature, vol. 518, no. 7539, pp. 360–364, Feb. 2015.

[25] B. D. Pope et al., “Topologically associating domains are stable units of replication-timing regulation,” Nature, vol. 515, no. 7527, pp. 402–405, Nov. 2014.

[26] M. S. Lawrence et al., “Mutational heterogeneity in cancer and the search for new cancer-associated genes,” Nature, vol. 499, no. 7457, pp. 214–218, Jul. 2013.

[27] H.-J. Wu and F. Michor, “A computational strategy to adjust for copy number in tumor Hi-C data,” Bioinformatics, vol. 32, no. 24, pp. 3695–3701, Dec. 2016.

[28] J. Dixon et al., “An Integrative Framework For Detecting Structural Variations In Cancer Genomes,” bioRxiv, p. 119651, Mar. 2017.

[29] D. Filippova, R. Patro, G. Duggal, and C. Kingsford, “Identification of alternative topological domains in chromatin,” Algorithms Mol. Biol., vol. 9, no. 1, p. 14, May 2014.

[30] C. Weinreb and B. J. Raphael, “Identification of hierarchical chromatin domains,” Bioinformatics, p. btv485, Aug. 2015.

[31] L. I. Malik and R. Patro, “Rich chromatin structure prediction from Hi-C data,” bioRxiv, p. 32953, Nov. 2015.

[32] C.-T. Ong and V. G. Corces, “CTCF: an architectural protein bridging genome topology and function,” Nat. Rev. Genet., vol. 15, no. 4, pp. 234–246, Apr. 2014.

[33] Z. Tang et al., “CTCF-Mediated Human 3D Genome Architecture Reveals Chromatin Topology for Transcription,” Cell.

[34] T.-H. S. Hsieh, A. Weiner, B. Lajoie, J. Dekker, N. Friedman, and O. J. Rando, “Mapping Nucleosome Resolution Chromosome Folding in Yeast by Micro-C,” Cell, vol. 162, no. 1, pp. 108–119, Jul. 2015.

[35] B. A. Bouwman and W. de Laat, “Getting the genome in shape: the formation of loops, domains and compartments,” Genome Biol., vol. 16, no. 1, p. 154, Aug. 2015.

[36] A. Harmanci, J. Rozowsky, and M. Gerstein, “MUSIC: identification of enriched regions in ChIP-Seq experiments using a mappability-corrected multiscale signal processing framework,” Genome Biol., vol. 15, no. 10, p. 474, Oct. 2014.

[37] F. Ay, T. L. Bailey, and W. S. Noble, “Statistical confidence estimation for Hi-C data reveals regulatory chromatin contacts,” Genome Res., vol. 24, no. 6, pp. 999–1011, Jun. 2014.

[38] M. Imakaev et al., “Iterative correction of Hi-C data reveals hallmarks of chromosome organization,” Nat. Methods, vol. 9, no. 10, pp. 999–1003, Oct. 2012.

[39] A. Fotuhi Siahpirani, F. Ay, and S. Roy, “A multi-task graph-clustering approach for chromosome conformation capture data sets identifies conserved modules of chromosomal interactions,” Genome Biol., vol. 17, p. 114, 2016.

[40] C. Dai et al., “Mining 3D genome structure populations identifies major factors governing the stability of regulatory communities,” Nat. Commun., vol. 7, p. 11549, May 2016.

[41] F. Ay and W. S. Noble, “Analysis methods for studying the 3D architecture of the genome,” Genome Biol., vol. 16, no. 1, p. 183, Sep. 2015.

[42] C. Lévy-Leduc, M. Delattre, T. Mary-Huard, and S. Robin, “Two-dimensional segmentation for analyzing Hi-C data,” Bioinformatics, vol. 30, no. 17, pp. i386–i392, Sep. 2014.

[43] J. Fraser et al., “Hierarchical folding and reorganization of chromosomes are linked to transcriptional changes in cellular differentiation,” Mol. Syst. Biol., vol. 11, no. 12, pp. 852–852, Dec. 2015.

[44] I. Rajapakse, D. Scalzo, S. J. Tapscott, S. T. Kosak, and M. Groudine, “Networking the nucleus,” Mol. Syst. Biol., vol. 6, no. 1, Jan. 2010.

[45] K. Kruse, S. Sewitz, and M. M. Babu, “A complex network framework for unbiased statistical analyses of DNA–DNA contact maps,” Nucleic Acids Res., vol. 41, no. 2, pp. 701–710, Jan. 2013.

[46] K.-K. Yan, D. Wang, J. Rozowsky, H. Zheng, C. Cheng, and M. Gerstein, “OrthoClust: an orthology-based network framework for clustering data across multiple species,” Genome Biol., vol. 15, no. 8, p. R100, Aug. 2014.

[47] N. Servant et al., “HiC-Pro: an optimized and flexible pipeline for Hi-C data processing,” Genome Biol., vol. 16, no. 1, p. 259, Dec. 2015.

[48] A. R. Barutcu et al., “Chromatin interaction analysis reveals changes in small chromosome and telomere clustering between epithelial and breast cancer cells,” Genome Biol., vol. 16, p. 214, 2015.

[49] K. C. Akdemir and L. Chin, “HiCPlotter integrates genomic data with interaction matrices,” Genome Biol., vol. 16, no. 1, p. 198, Sep. 2015.

[50] E. Crane et al., “Condensin-driven remodelling of X chromosome topology during dosage compensation,” Nature, vol. 523, no. 7559, pp. 240–244, Jul. 2015.

[51] R. Nussinov, G. Pieczenik, J. Griggs, and D. Kleitman, “Algorithms for Loop Matchings,” SIAM J. Appl. Math., vol. 35, no. 1, pp. 68–82, Jul. 1978.

